# Next-Generation Imipridones ONC206 and ONC212 Synergize with Lurbinectedin in Killing Pancreatic Ductal Adenocarcinoma Cells

**DOI:** 10.64898/2026.08.13.744614

**Authors:** Tej Tummala, Audrey Su, Ashley Sanchez Sevilla Uruchurtu, Christopher G. Azzoli, Wafik S. El-Deiry

**Affiliations:** Laboratory of Translational Oncology and Experimental Cancer Therapeutics, Warren Alpert Medical School, Brown University, Providence, RI 02903, USA; Legorreta Cancer Center at Brown University, Providence, RI 02912, USA; Icahn School of Medicine at Mount Sinai, New York, NY, USA; Department of Pathology and Laboratory Medicine, Warren Alpert Medical School, Brown University, Providence, RI 02903, USA; Joint Program in Cancer Biology, Brown University Health System and Brown University, Providence, RI 02903, USA; Hematology/Oncology Division, Department of Medicine, Brown University Health System and Brown University, Providence, RI 02903, USA

**Keywords:** PDAC, lurbinectedin, ONC206, ONC212, synergy

## Abstract

Pancreatic ductal adenocarcinoma (PDAC) is a devastating malignancy with a five-year survival rate of approximately 13%, underscoring the urgent need for novel therapeutic strategies. Next-generation imipridones ONC206 and ONC212 are potent anticancer agents that activate the mitochondrial ClpP protease and the integrated stress response. Lurbinectedin, an FDA-approved therapy for metastatic small cell lung cancer, inhibits transcription by binding the DNA minor groove and has demonstrated preclinical efficacy in PDAC models. Here, we show that ONC206 and ONC212 are highly cytotoxic against PDAC cell lines as monotherapies and in combination with lurbinectedin. Both ONC206 and ONC212 achieved sub-micromolar seventy-two-hour IC₅₀ values in BxPC-3, PANC-1, and HPAF-II PDAC cells, with ONC212 exhibiting greater potency across all lines. Mechanistically, ONC206 and ONC212 induce apoptosis through ClpX depletion, ATF4 induction, and caspase-mediated PARP cleavage. Combination treatment of lurbinectedin with both imipridones produced robust synergy, with ONC212 generally exhibiting stronger synergy at lower concentrations and HSA synergy scores up to 29.5. Importantly, these combinations showed minimal toxicity in CCD 841 CoN non-malignant colon epithelial cells, indicating selective tumor cell killing. Western blot analysis revealed that synergy between lurbinectedin and ONC212 is associated with upregulation of DR5 and downregulation of Bcl-2 and ClpX. These findings provide mechanistic and preclinical support for combining lurbinectedin with next-generation imipridones as a therapeutic strategy in PDAC.

## Introduction

Pancreatic cancer remains a devastating disease with a five-year survival rate of approximately 13% [1]. While surgical resection provides the best chance at long-term survival, only 15-20% of pancreatic cancer is resectable at the time of diagnosis [2]. Pancreatic ductal adenocarcinoma (PDAC) accounts for 90% of pancreatic cancer cases, making it the most prevalent pancreatic neoplasm [3]. Despite novel research into the tumor biology of PDAC and advancements in treatments, current therapies have showed limited success, highlighting the need for novel anti-tumor therapies.

ONC212 and ONC206 are next-generation imipridones and analogues of ONC201, a first-in-class imipridone, with promising preclinical activity against a variety of tumor types, including pancreatic cancer, through integrated stress response and mitochondrial pathways. ONC212 is a fluorinated imipridone that binds and suppresses ClpX, the regulatory binding partner of ClpP, impairing oxidative phosphorylation (OXPHOS), decreasing mitochondrial-derived ATP production, and inducing apoptosis in OXPHOS-dependent cells [4, 5]. Additionally, ONC212 has been shown to induce caspase-3 activation, leading to apoptosis in a mitochondrial-independent manner [6]. ONC212 has shown activity against pancreatic cancer as a preclinical monotherapy or combinatorial therapy with 5-fluoruacil, irinotecan, or oxaliplatin [7]. ONC206, similarly, has been shown to induce the integrated stress response, the TRAIL pathway, and ClpP activation, leading to mitochondrial damage [8]. In general, ONC206 and ONC212 have shown greater anti-tumor activity than ONC201, with ONC212 demonstrating efficacy in ONC201-resistant solid tumors and ONC206 demonstrating higher potency than ONC201 in killing a variety of solid tumors [7, 9–11].

Lurbinectedin (Zepzelca, PM01183) is a tetrahydropyrroloquinoline and synthetic analog of trabectedin that covalently binds to CG-rich regions in the minor groove of DNA, leading to stalling and degradation of RNA Polymerase II [12, 13]. After forming DNA adducts, lurbinectedin has been shown to induce double-stranded DNA breaks, triggering S-phase accumulation and apoptosis [13–15]. Further, lurbinectedin has been shown to impact the tumor microenvironment through a reduction of tumor-associated macrophages and circulating monocytes, while also inhibiting the production of inflammatory and angiogenic factors CCL2, CXCL8, and VEGF [16].

Lurbinectedin was FDA approved in 2020 for the treatment of adults with metastatic small cell lung cancer (SCLC) with disease progression on or after platinum-based chemotherapy [13, 17, 18]. Currently, it is undergoing clinical trials in a variety of tumor types. The first human phase I study of lurbinectedin began in 2009 in patients with a variety of advanced solid tumors and found partial response in a patient with pancreatic adenocarcinoma [19]. A pooled safety analysis of lurbinectedin in 554 patients with a variety of advanced solid tumors, primarily ovarian cancers, SCLC, and endometrial cancers, concluded lurbinectedin has a manageable safety profile in advanced solid tumors [20].

Preclinical data supports lurbinectedin’s candidacy in pancreatic adenocarcinoma. Lurbinectedin was recently shown to exhibit sub-nanomolar seventy-two-hour IC₅₀ in PDAC cell lines HPAF-II, PANC-1, and BxPC-3. Further, it was shown to synergize with chemotherapeutics irinotecan and 5-fluorouracil in killing PDAC cell lines in vitro [21]. Lurbinectedin has also been shown to synergize with gemcitabine in PDAC mouse models [22]. Finally, lurbinectedin has been shown to synergize with first-generation imipridone ONC201, inducing potent cytotoxicity at low concentrations in SCLC cell lines [23].

Our group explored the potency of imipridones ONC206 and ONC212 in pancreatic adenocarcinoma cell lines as single agents and in combination with lurbinectedin. We report highly efficient killing of PDAC cell lines by both ONC206 and ONC212 monotherapy and potent synergism in conjunction with lurbinectedin without excessive toxicity in regular colon epithelial cells.

## Methods

### Cell Lines and Culture Conditions

Human pancreatic cancer cell lines PANC-1 (CRL-1469), HPAF-II (CRL-1997), and BxPC-3 (CRL-1687) were procured from the American Type Culture Collection (ATCC). PANC-1 was derived from a pancreatic ductal tumor of a 56-year-old Caucasian male diagnosed with epithelioid carcinoma. HPAF-II originated from a 44-year-old male patient with metastatic pancreatic adenocarcinoma. BxPC-3 was isolated from the tissue of a 61-year-old female with pancreatic adenocarcinoma. The KRAS gene is mutated in both PANC-1 and HPAF-II, whereas BxPC-3 retains a wild-type KRAS allele. All cell lines harbor TP53 mutations [24]. Cells were cultured in Dulbecco’s Modified Eagle Medium (DMEM) supplemented with 10% fetal bovine serum (FBS) and 1% penicillin–streptomycin. Standard incubation conditions of 37 °C in a humidified environment with 5% CO₂ were maintained throughout the experiments.

### Therapeutic Compounds

Lurbinectedin (MedChem Express, Cat# HY-16293), ONC206 (Chimerix), and ONC212 (Chimerix) were used as therapeutic agents. All compounds were stored at −20 °C in accordance with the manufacturer’s guidelines until use.

### Cell Viability and Synergy Analysis

Cell viability was assessed using the CellTiter-Glo (CTG) luminescence assay. Approximately 3,000 cells were seeded per well into black 96-well plates and allowed to adhere overnight. After 24 hours, cells were treated with increasing concentrations of lurbinectedin, ONC206, or ONC212 to determine individual IC₅₀ values. Drug dilutions were prepared via 1:2 serial dilution, and plates were incubated for seventy-two hours post-treatment. Viability was determined by measuring ATP levels via luminescence, which correlates with metabolic activity. Luminescence values were normalized to untreated control wells to calculate the percentage of viable cells.

To evaluate combinatorial effects, dual-drug treatments were carried out using the same seeding and incubation conditions. Lurbinectedin and ONC206 or ONC212 were applied at varying concentrations across the plate, with the upper leftmost well serving as the no-drug control. Following seventy-two hours of treatment, CTG reagent was added, and luminescence was recorded. Normalized viability data were subjected to synergy analysis using the SynergyFinder software platform, employing the Highest Single Agent (HSA) reference model. The HSA synergy score was calculated using the equation SHSA = EA,B,…,N − max(EA, EB,…, EN), where EA,B,…,N represents the effect of the drug combination and EA, EB,…, EN are the responses to individual agents. A synergy score >10 denotes a strongly synergistic interaction, <−10 indicates antagonism, and scores between −10 and 10 suggest additivity [25, 26].

### Western Blotting

Western blotting was conducted to assess protein expression changes in response to single and combination drug treatments in PANC-1, BxPC-3, and HPAF-II cells. Cells were plated at a density of ∼500,000 cells per well in 6-well plates and treated with vehicle or drug(s) after overnight adherence. After 48 hours of treatment, cells were harvested using a cell scraper, lysed in RIPA buffer supplemented with protease and phosphatase inhibitors, and subjected to protein quantification via a BCA assay (Pierce BCA Kit, Life Technologies, Cat# 23225). Equal amounts of protein were separated on 4–12% SDS-PAGE gels and transferred to PVDF membranes. Membranes were blocked in 5% non-fat dry milk in TBST for 30 minutes before overnight incubation at 4 °C with primary antibodies diluted in blocking buffer, according to manufacturer-recommended concentrations. Secondary antibodies—Goat anti-Rabbit IgG (HRP, Invitrogen Cat# 31460) and Goat anti-Mouse IgG (HRP, Invitrogen Cat# 31430)—were used at 1:5000 dilution. Membranes were developed using enhanced chemiluminescence (ECL).

## Results

### ONC212 exhibits greater cytotoxicity than ONC206 in PDAC cell lines at low concentrations

Both ONC212 and ONC206 exhibited sub-micromolar seventy-two-hour IC₅₀ values in all three PDAC cell lines (HPAF-II, PANC-1, BxPC-3). ONC212 demonstrated markedly lower values IC₅₀ across each PDAC cell line compared to ONC206, indicating that ONC212 is the more potent imipridone under these conditions (**Figure 1**).

**Figure 1.**
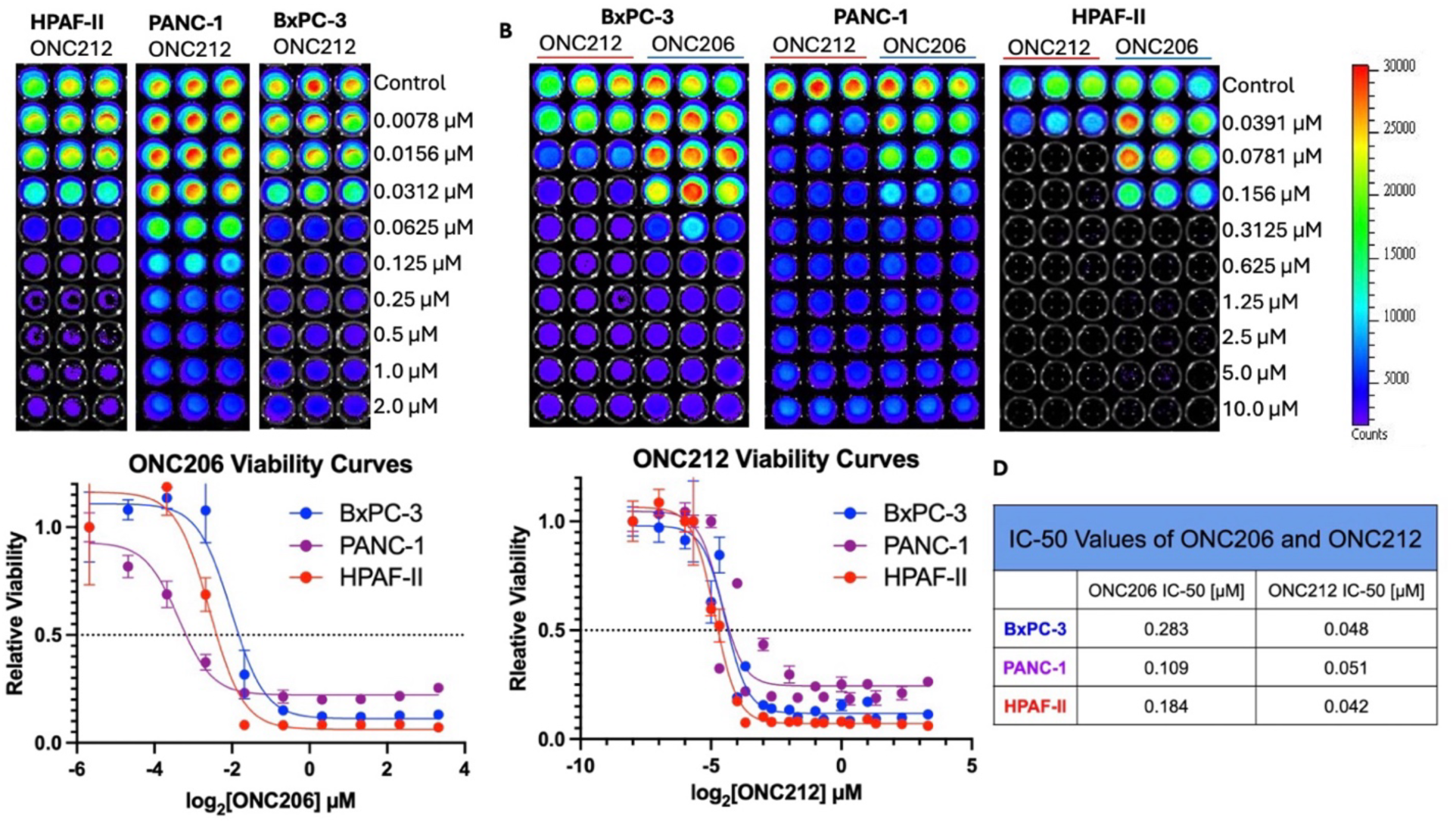
ONC212 is a more potent cytotoxic agent than ONC206 in killing PDAC cell lines over seventy-two hours of treatment. HPAF-II, PANC-1, and BxPC-3 were treated with low to sub-micromolar concentrations of ONC212 (A and B) or ONC206 (B) monotherapy. Relative cell-viability was normalized to the control and plotted (C) and seventy-two-hour IC₅₀ values were calculated and displayed (D).

The sub-micromolar seventy-two-hour IC₅₀ values observed for both ONC206 and ONC212 across the three PDAC cell lines motivated further investigation into each drug’s mechanism of action. To better understand the molecular basis of cell death, western blots were performed to analyze ONC206 and ONC212 treatment across multiple timepoints and concentrations in the PANC-1 cell line.

### ONC206 and ONC212 induce apoptosis through mitochondrial and integrated stress response pathways in human PANC1 pancreatic adenocarcinoma cells

We investigated the effects of ONC206 and ONC212 on depletion of mitochondrial ClpX as a marker of mitochondrial ClpP target engagement, and upregulation of ATF4 as a marker of integrated stress response activation (**Figure 2**). Both ONC206 and ONC212 displayed a dose-dependent suppression of ClpX expression and induction of ATF4, with ONC212 showing the effects at lower doses consistent with its higher potency. The effects were observed at early and late time points with the later time points showing PARP cleavage indicative of caspase-dependent induction of apoptosis (**Figure 2**).

**Figure 2.**
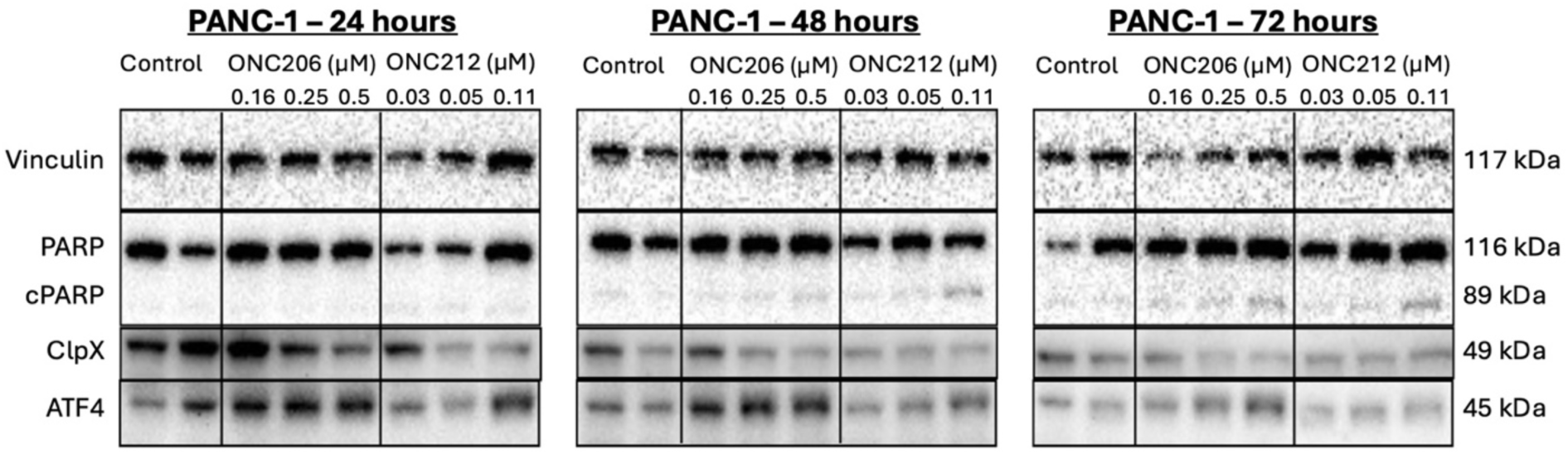
ONC206 and ONC212 involve mitochondrial and integrated stress response pathways to induce apoptosis of PANC-1 cells. The panels show western blots of effects of either ONC206 or ONC212 at different doses and time points compared with controls as indicated. cPARP, a measure of caspase-dependent apoptosis, increases in a time- and concentration-dependent manner in both ONC206 and ONC212 (as indicated). ONC212 shows greater potency at lower concentrations and earlier timepoints. Across all timepoints, ONC206 and ONC212 decrease ClpX levels. In 24 and 48 hours, ONC206 and ONC212 upregulate ATF4 in a concentration-independent and concentration-dependent manner, respectively.

### Lurbinectedin exhibits potent synergy in combination with ONC206 and ONC212 in killing human BxPC-3 PDAC cells with minimal toxicity to benign colon epithelial cells

Given the potent single-agent activity of both imipridones, ONC206 and ONC212 were tested in combination with lurbinectedin to determine if combinatorial therapy would result in synergistic killing of tumor cells (**Figure 3**). The results indicate that both ONC206 and ONC212 demonstrate potent synergy when combined with lurbinectedin at multiple dose combinations (**Figure 3**).

**Figure 3.**
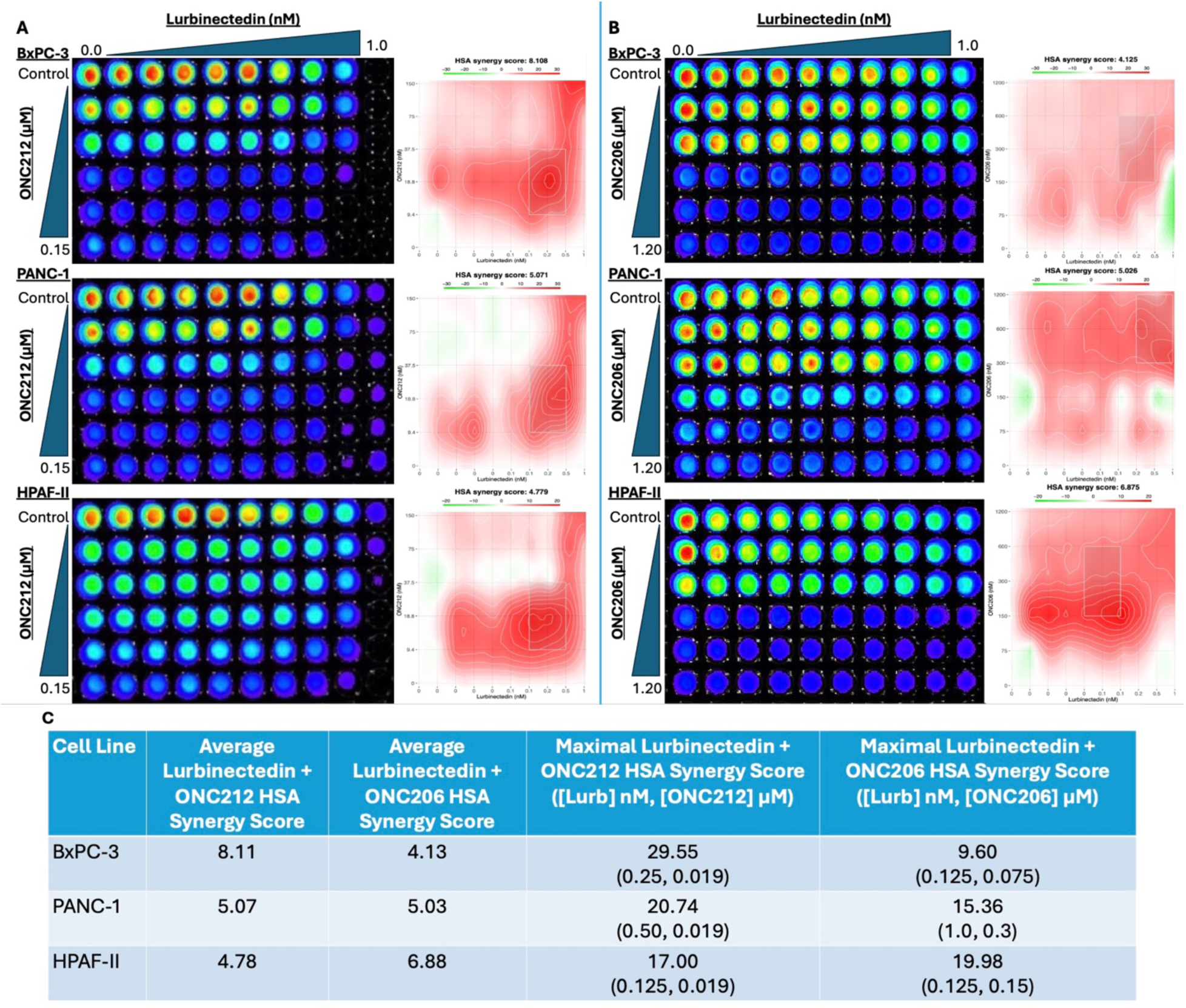
Both ONC212 and ONC206 synergize with lurbinectedin in killing human PDAC cell lines after seventy-two hours of treatment. Across each cell line shown in panels A (ONC212 and lurbinectedin) and B (ONC206 and lurbinectedin), there were consistently positive HSA synergy scores (C), demonstrating synergism between lurbinectedin and the imipridones across all three cell lines BcPC-3, PANC-1, and HPAF-II. Generally, ONC212 with lurbinectedin showed stronger or comparable synergism compared to ONC206 with lurbinectedin, yet lurbinectedin with ONC206 seemed more potent in HPAF-II cells. The highest overall synergism was observed in lurbinectedin with ONC212 with an HSA score of 29.55. HSA scores > 10 are often deemed strongly synergistic.

Once potent synergism between the next-generation imipridones and lurbinectedin was established in PDAC cell lines, the combinatorial therapies were tested in CCD 841 CoN healthy colon epithelial cells to determine if there was excessive toxicity to non-cancerous cells (**Figure 4**).

**Figure 4.**
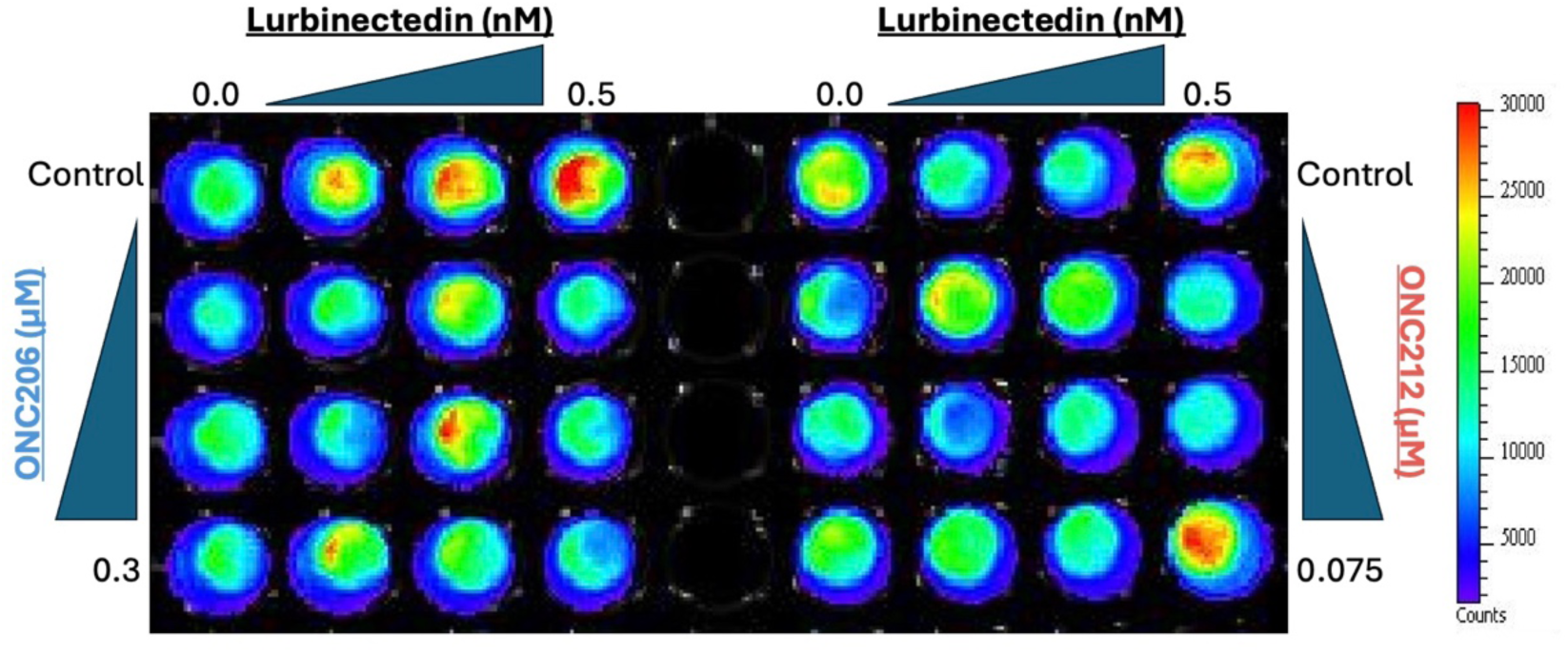
Combinatorial therapies of lurbinectedin with ONC206 and ONC212 both demonstrate minimal killing of CCD 841 CoN healthy human colon epithelial cells after 48 hours of treatment. CTG imaging shows high viability of healthy colon epithelial cells at dosages of lurbinectedin with either ONC206 or ONC212 that induced significant cytotoxicity in PDAC cell lines.

### The combination of lurbinectedin with ONC212 suppresses Bcl2 and increases DR5 expression in human PDAC cells

After determining minimal cytotoxicity to healthy colon epithelial cells, the combination of lurbinectedin with ONC212 was subjected to western blot analysis to better elucidate the mechanisms underlying synergism (**Figure 5**). The suppression of Bcl2 was more evident in HPAF-II cells as compared with BxPC-3 after either ONC212 monotherapy or in combination with lurbinectedin. TRAIL receptor DR5 was increased by ONC212 in both BxPC-3 and HPAF-II including when it was combined with lurbinectedin. ClpX expression was suppressed by ONC212 monotherapy or in combination with lurbinectedin indicating target engagement with mitochondrial ClpP in both BxPC-3 and HPAF-II cells (**Figure 5**).

**Figure 5.**
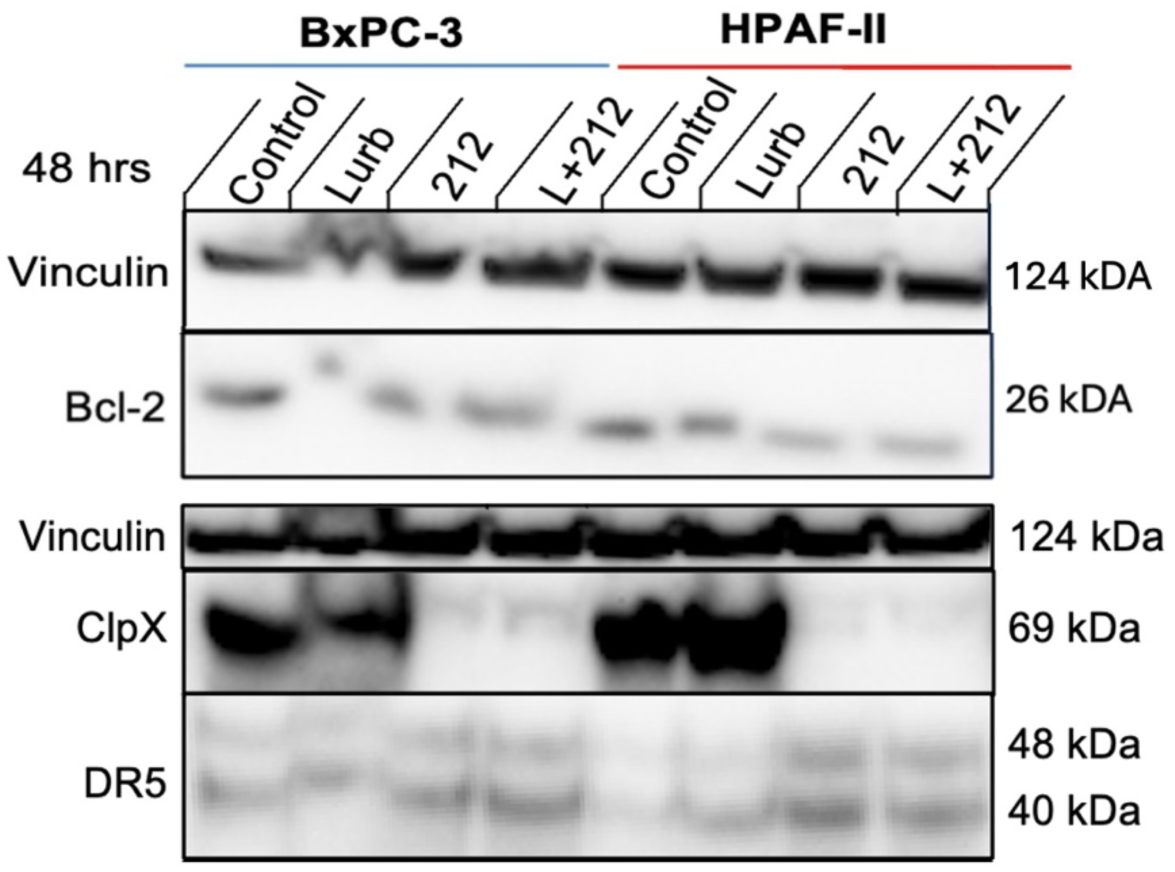
Effects of lurbinectedin and ONC212 on apoptotic and mitochondrial stress markers in PDAC cell lines at 48 hours. Western blot analysis was conducted on HPAF-II and BxPC-3 cells after exposure to lurbinectedin and ONC212 monotherapies and combinatorial therapy. In both cell lines, ONC212 markedly reduced ClpX levels, indicating activation of mitochondrial protease ClpP, with lurbinectedin contributing minimally to ClpX expression. Bcl-2, an anti-apoptotic protein, appears decreased by combinatorial therapy, demonstrating a suppression of anti-apoptotic signaling. DR5 expression seems slightly elevated in both ONC212 and lurbinectedin monotherapies and combinatorial therapy, indicating activation of stress-induced apoptotic pathways. Vinculin was used as a protein loading control.

## Discussion

Pancreatic ductal adenocarcinoma (PDAC) remains one of the most lethal solid malignancies. This study demonstrates that next generation imipridones ONC206 and ONC212 exhibit potent cytotoxicity in PDAC cell lines as monotherapies or in combination with lurbinectedin. Our results provide insights into novel combinatorial regimens and elucidating the molecular mechanisms underlying tumor cell cytotoxicity.

Across PDAC cell lines BxPC-3, PANC-1, and HPAF-II, both ONC206 and ONC212 demonstrated sub-micromolar IC₅₀ values, yet ONC212 was significantly more potent than ONC206 (Figure 1). These results are consistent with prior research comparing ONC201, ONC206, and ONC212. Borsuk et al. and Chang et al. found ONC212 to be the most potent imipridone followed by ONC206 and ONC201 in a variety of tumor types [27, 28]. Similarly, Wagner et al. demonstrated ONC206 and ONC212 demonstrate improved anti-cancer activity compared to ONC201 without excessive toxicity to human fibroblasts [29].

Timepoint western blot analysis of ONC206 and ONC212 in PANC-1 cells indicate apoptosis is mediated by caspase-mediated PARP cleavage, depletion of ClpX, and activation of the integrated stress response through upregulation of ATF4 (Figure 2). Again, these findings are consistent with previous mechanistic studies of imipridones revealing they frequently induce mitochondrial dysfunction and stress-mediated apoptosis [4, 6, 7]. cPARP is a measure of apoptotic cell death, and ONC212 induces similar levels of PARP cleavage in comparison to ONC206 at markedly lower concentrations, further supporting that ONC212 is a more potent cytotoxic agent than ONC206 in PDAC cell lines [30].

Given the potency of ONC206 and ONC212 monotherapies, they were combined with lurbinectedin, and synergism was observed across all tested PDAC cell lines at sub-nanomolar and nanomolar concentrations of lurbinectedin and imipridones, respectively (Figure 3). HSA scores exceeded 10 in each combinatorial therapy, denoting strong synergism in accordance with HSA synergy criteria [25, 26]. In general, lurbinectedin with ONC212 seemed to achieve more robust synergism, with the highest average synergy score occurring in the lurbinectedin-ONC212 combination in BxPC-3 cell lines, with scores exceeding 29.5, indicative of strong synergism. However, lurbinectedin with ONC206 was, on average, more synergistic in HPAF-II cells, indicating that lurbinectedin-imipridone synergy is cell-line-specific.

Importantly, combinatorial therapies of lurbinectedin with imipridones demonstrated minimal cytotoxicity in non-malignant CCD 841 CoN colon epithelial cells (Figure 4). The selective killing of tumor cells under combinatorial therapy suggests a favorable therapeutic index which likely reflects the enhanced dependence of PDAC cells on transcription, mitochondrial function, and stress-response pathways, which are inhibited by lurbinectedin and imipridones.

Due to the greater potency of ONC212 monotherapy in PDAC compared to ONC206 monotherapy, generally stronger synergism between lurbinectedin with ONC212 over lurbinectedin with ONC206, and comparable minimal toxicity to healthy colon epithelial cells, lurbinectedin and ONC212 treated BxPC-3 and HPAF-II cells were subjected to western blot to analyze the cellular mechanisms underlying the combinatorial therapy. In both cell lines, combinatorial therapy indicated depletion of ClpX, downregulation of Bcl-2, and upregulation of DR5 (Figure 5). ClpX depletion is consistent with uncontrolled ClpP activation, contributing to stress-response activation and downstream apoptosis [4]. DR5, also known as TRAIL-R2, is a member of the tumor necrosis factor receptor superfamily, and upon binding to TRAIL (TNF-related inducing ligand), will lead to apoptosis [31]. DR5 upregulation is consistent with previous findings that imipridones modulate cell death via induction of the TRAIL pathway [8]. Bcl-2, an anti-apoptotic protein that inhibits mitochondrial apoptosis, decreases under combinatorial therapy, providing another mechanism for cytotoxicity [32]. It is plausible that lurbinectedin’s induction of DNA damage and transcriptional inhibition sensitizes PDAC cell lines to imipridone-induced mitochondrial stress, enabling potent synergistic killing of PDAC cells at low drug concentrations, although more mechanistic data would be needed to validate such a hypothesis [14].

Collectively, these results expand on previous work that demonstrates lurbinectedin synergizes with ONC201 in small cell lung cancer [23]. Further, it builds on previous research indicating that lurbinectedin is a potent cytotoxic agent as both a monotherapy and in combination with gemcitabine, irinotecan, and 5-fluorouracil in PDAC models [21, 22].

Our findings strongly suggest that ONC206 and ONC212 are potent anti-cancer agents in PDAC cell lines as monotherapies or in combination with lurbinectedin. Mitochondrial and integrated stress response pathways are critical for imipridone-induced apoptosis. Similarly, ONC212 and lurbinectedin’s potent synergism at nanomolar and sub-nanomolar concentrations, respectively, is mediated by mitochondrial stress, the integrated stress response, and the TRAIL-DR5 apoptotic pathway, without causing excessive toxicity to healthy colon epithelial cells. Our findings provide preclinical evidence supporting the therapeutic potential of combining lurbinectedin with next-generation imipridones to treat PDAC.

## Conclusion

Our findings demonstrate that next-generation imipridones ONC206 and ONC212 exhibit potent cytotoxic activity against PDAC cell lines BxPC-3, PANC-1, and HPAF-II as monotherapies or in combination with lurbinectedin without excessive toxicity to non-malignant epithelial cells. Mechanistic analysis shows that mitochondrial stress, integrated stress response signaling, and apoptotic pathways all contribute to potent drug synergism at nanomolar and sub-nanomolar concentrations of imipridones and lurbinectedin, respectively. These results provide preclinical support for a combinatorial therapy of lurbinectedin with imipridones, or imipridone monotherapy, to treat PDAC.

Future studies should evaluate and compare these combinatorial therapies in in vivo PDAC models. Further, future mechanistic studies should further analyze mitochondrial function, integrated stress response pathways, DNA damage, transcription inhibition, and various apoptotic pathways to better understand cellular mechanisms behind synergism.

## Acknowledgements

W.S.E-D. is an American Cancer Society Research Professor and is supported by the Mencoff Family University Professorship at Brown University. This work was supported by an NIH grant (CA173453) to W.S.E-D. This work was presented in part at the 2023 (Tummala, T., Sanchez Sevilla Uruchurtu, A., *et al*. Synergistic Combinations of Lurbinectedin with Irinotecan and ONC212 in Pancreatic Cancer. 114th Annual AACR meeting, April 14-19, 2023, Orlando, FL.), 2024 (Tummala, T., Sevilla Uruchurtu, A.S., *et al*. Synergistic combinations of lurbinectedin with ONC212 in pancreatic cancer. 115th Annual AACR meeting, April 5-10, 2024, San Diego, CA.), 2025 (Tummala, T., Uruchurtu, A.A.S., *et al*. Preclinical analysis of ONC206 and ONC212 with lurbinectedin in pancreatic cancer. 116th Annual AACR meeting, April 25-30, 2025, Chicago, IL.) meeting of the American Association for Cancer Research.

